# Molecular dating for phylogenies containing a mix of populations and species

**DOI:** 10.1101/536656

**Authors:** Beatriz Mello, Qiqing Tao, Sudhir Kumar

## Abstract

Concurrent molecular dating of population and species divergences is essential in many biological investigations, including phylogeography, phylodynamics, and species delimitation studies. Multiple sequence alignments used in these investigations frequently consist of both intra- and inter-species samples (mixed samples). As a result, the phylogenetic trees contain inter-species, inter-population, and within population divergences. To date these sequence divergences, Bayesian relaxed clock methods are often employed, but they assume the same tree prior for both inter- and intra-species branching processes and require specification of a clock model for branch rates (independent vs. autocorrelated rates models). We evaluated the impact of using the same tree prior on the Bayesian divergence time estimates by analyzing computer-simulated datasets. We also examined the effect of the assumption of independence of evolutionary rate variation among branches when the branch rates are autocorrelated. Bayesian approach with Skyline-coalescent tree priors generally produced excellent molecular dates, with some tree priors (e.g., Yule) performing the best when evolutionary rates were autocorrelated, and lineage sorting was incomplete. We compared the performance of the Bayesian approach with a non-Bayesian, the RelTime method, which does not require specification of a tree prior or selection of a clock model. We found that RelTime performed as well as the Bayesian approach, and when the clock model was mis-specified, RelTime performed slightly better. These results suggest that the computationally efficient RelTime approach is also suitable to analyze datasets containing both populations and species variation.

## Introduction

Divergence times derived from molecular data have become critical for elucidating earth’s historical processes that have shaped the evolution of life (Hedges, Marin, Suleski, Paymer, & Kumar, 2015; Ho, 2014; O’Reilly, dos Reis, & Donoghue, 2015). Recent biological timescales of closely-related taxa are generally built in the context of phylogeographic, phylodynamics and species delimitation studies (Esselstyn, Evans, Sedlock, Anwarali Khan, & Heaney, 2012; McCluskey & Postlethwait, 2015; Mello, Vilela, & Schrago, 2018; C. Wang, Shikano, Persat, & Merilä, 2015). Divergence times are usually estimated from molecular data with multiple individuals per species sampled. Consequently, both inter- and intra-species divergences (nodes) are present in the same phylogenetic tree, such that both speciation and population processes are at play. Such datasets are becoming increasingly common (Edwards, Shultz, & Campbell-Staton, 2015; Lemmon & Lemmon, 2012; Manzanilla et al., 2018; McCormack, Hird, Zellmer, Carstens, & Brumfield, 2013; Melville et al., 2017; Merwe, McPherson, Siow, & Rossetto, 2014).

In order to accurately infer divergence times in datasets with a mixture of micro- and macro-evolutionary events, a multispecies coalescent (MSC) approach would be highly appropriate, because it explicitly accounts for conflicts between gene genealogies and the species tree by modelling incomplete lineage sorting (ILS) across lineages (Degnan & Salter, 2005; Edwards & Beerli, 2000; Heled & Drummond, 2010; Maddison, 1997; Rannala & Yang, 2003; Takahata, 1989; Takahata & Nei, 1985; Yang & Rannala, 2014). However, MSC is not often applied because computational times required to estimate divergence times are still impractical for contemporary data sizes, even though some recent advances have been made to reduce the computational burden (Ogilvie, Bouckaert, & Drummond, 2017; Ogilvie, Heled, Xie, & Drummond, 2016; Rannala & Yang, 2017; Xu & Yang, 2016). Moreover, it currently requires *a priori* delimitation of species limits and, for some implementations, the assumption of strict molecular clocks (Heled & Drummond, 2010; Xu & Yang, 2016; Yang, 2015).

Instead, researchers frequently use the standard Bayesian framework, mainly the BEAST software (Bouckaert et al., 2014; dos Reis & Yang, 2011). BEAST software requires specification of priors, including models that describe the branching pattern (tree prior) and that assume the absence of autocorrelated branch rates (ABR). The commonly used tree priors in BEAST analyses are those that describe speciation processes, e.g., Yule (Yule, 1924) and birth and death (BD) (Feller, 1939; Kendall, 1948), and within-species population processes, e.g., coalescent priors considering a constant population size or an exponential growth (Kuhner, Yamato, & Felsenstein, 1995, 1998). Yule and BD priors model macro-evolutionary events, so their use to describe intra-species divergences would improperly assume these divergences to be like speciation events rather than due to intra-population coalescence. On the other hand, the use of coalescent priors may bias time estimation of ancient inter-species divergences, like the tMRCA (time to the most recent common ancestor) of a whole genus or several extant species.

Currently, in the standard Bayesian molecular dating, there is no composite tree prior that account for macro- and micro-evolutionary processes in estimating divergence times. Therefore, it is critical to investigate whether the divergence times produced by Bayesian methods are robust to the selection of tree prior and to the application of only one tree prior for trees with mixed samples. Also, the relaxed clock method in BEAST assumes that there is no autocorrelation of evolutionary rates among branches. However, recent evidence suggests that autocorrelated branch rates are likely to be the norm (Tao, Tamura, Battistuzzi, & Kumar, 2019), so the impact of rate model misspecification in using BEAST needs to be evaluated.

Ritchie et al. (2017) have already conducted some simulations to evaluate the impact of tree priors on time estimates in BEAST. In their study, they only considered the case where data were simulated under an independent branch rate (IBR) model and with incomplete lineage sorting (ILS) (**Table 1**). Therefore, we do not yet know the impact of using a single tree prior on time estimates in BEAST under many other scenarios, such as an ABR model and the absence of ILS. Furthermore, Ritchie et al. (2017) did not evaluate the performance of any non-Bayesian method. For example, RelTime has been shown to work well for trees consisting exclusively of interspecies divergences (Mello, Tao, Tamura, & Kumar, 2017; Tamura, Tao, & Kumar, 2018), so its absolute and comparative performance will inform about its suitability for use in datasets containing both inter- and intra-species sampling. Thus, we conducted many simulated analyses that present a significant advance beyond Ritchie et al. (2017) (**Table 1**). They uniquely provide insights into many questions that are faced by practitioners in molecular dating studies.

**Table 1.** Computer simulation scenarios considered in this article.

|  | BEAST | BEAST | RelTime | RelTime |
| --- | --- | --- | --- | --- |
| Branch rates | ILS | No ILS | ILS | No ILS |
| Constant (CBR) | New | New | New | New |
| Independent (IBR) | Ritchie et al. | New | New | New |
| Autocorrelated (ABR) | New | New | New | new |

ILS, the presence of incomplete lineage sorting. No-ILS, the absence of incomplete lineage sorting.

## Material and methods

We analyzed simulated datasets to test the performance of a Bayesian approach (BEAST) (Drummond, Ho, Phillips, & Rambaut, 2006) and RelTime method (Kumar, Stecher, Li, Knyaz, & Tamura, 2018; Tamura et al., 2012). Simulations provide valuable means to test the reliability of inferred node ages because the actual time is known, and the impact of many biologically realistic conditions can be explored, including the presence/absence of autocorrelation of branch rates across the tree (ABR vs. IBR model) and incomplete lineage sorting (ILS vs. no-ILS) (**Fig. 1**).

**Figure 1.**
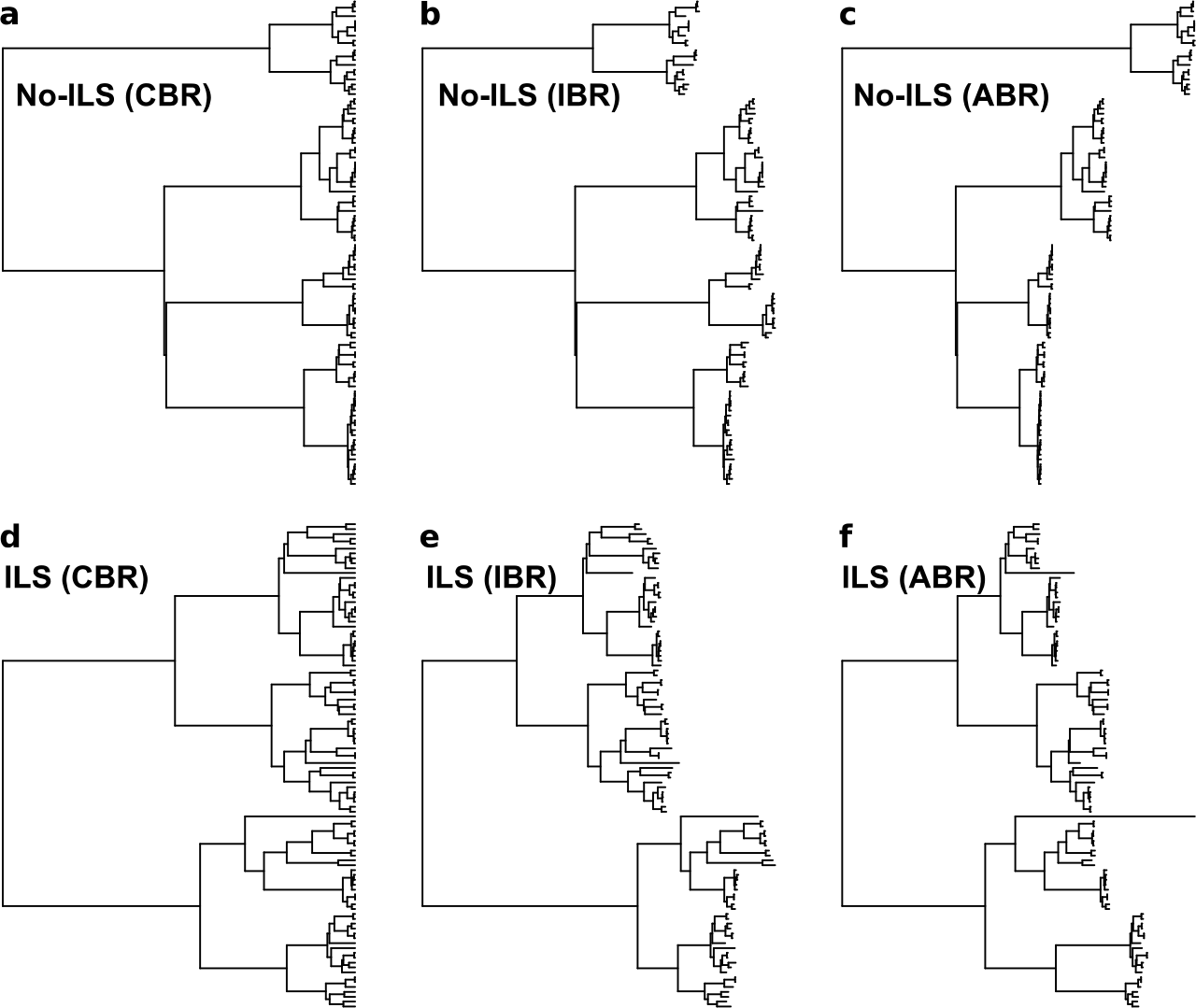
An example of the diversity of trees used in this study. Panels **a-c**, trees simulated without lineage sorting (no-ILS); **d-e**, trees simulated with incomplete lineage sorting (ILS). **a** and **d** were simulated under a constant branch rate (CBR) model; **b** and **e** were simulated under an independent branch rate (IBR) model; and **c** and **f** were simulated under an autocorrelated branch rate (ABR) model.

### Computer Simulated Datasets

We considered two distinct scenarios to simulate datasets with a mixed sampling of intra- and inter-species sequences. First, we assumed that there was complete lineage sorting (no-ILS), which represents the baseline scenario for understanding sources of error. In no-ILS phylogenies, all species are reciprocally monophyletic and speciation times are concordant with the divergence of sequences (e.g., **Fig. 1a-c**). Second, genealogies were simulated within species phylogeny under the MSC approach, which resulted in ILS. In this case, the sequence phylogeny is not fully concordant with the species tree, and species are not automatically monophyletic (e.g., **Fig. 1d-f**). The ILS scenario allowed us to assess the performance of methods under more complex evolutionary histories.

The no-ILS phylogenies contained ten species and were simulated assuming a Yule process with λ = 0.11 (speciation rate) and a root height of 10 million years (Ma) in the TreeSim R package (Stadler, 2011). Then, the *ms* function from phyclust R package, which generates coalescent trees under a modified Hudson (2002)’s neutral model, was used to simulate coalescent trees under constant population size with ten samples (*k* = 10) (Chen, 2011; Hudson, 2002). Assumed generation time was 20 years, and effective population size (*N*_*e*_) was 50,000. Coalescent trees were then “pasted” at each tip (species) of the phylogeny while keeping the expected phylogeny ultrametric because all the sequences were contemporaneous. For phylogenies with ILS, the same procedure and parameters were used to estimate species phylogenies (Yule process with ten species, λ = 0.11, and root height = 10 Ma). Then, single gene genealogies were simulated in Phybase R package (Liu & Yu, 2010) with a constant scaled population parameter (theta) of 0.014 and ten individual samples per species, which is equivalent to the configuration used to simulate coalescent trees in the scenario without ILS. An evolutionary rate of 3.5×10^−9^ substitutions/site/year (Endicott & Ho, 2008) was used, and one hundred datasets were obtained for both ILS and no-ILS analysis.

We simulated three different types of evolutionary rate variation among branches: constant branch rates (CBR, **Fig. 1a** and **d**), independent branch rates (IBR, **Fig. 1b** and **e**), and autocorrelated branch rates (ABR, **Fig. 1c** and **f**). In CBR, the evolutionary rate of 3.5×10^−9^ substitutions/site/year was applied to all branches of the phylogenies. In IBR, branch rates were allowed to randomly vary as much as fifty percent of the mean (Tamura et al., 2012). In ABR, we used modified functions from NELSI R package (Ho, Duchêne, & Duchêne, 2015) using 3.5×10^−9^ substitutions/site/year as the initial rate and a correlation parameter (*v*) of 0.01 (Kishino, Thorne, & Bruno, 2001). Sequences were simulated in seq-gen (Rambaut & Grass, 1997) under HKY substitution model (Hasegawa, Kishino, & Yano, 1985) to generate alignments of 10,000 sites with equilibrium frequencies and transition/transversion ratios sampled from an empirical distribution (Rosenberg & Kumar, 2003). Thus, we simulated six scenarios: CBR – no-ILS, IBR – no-ILS, ABR – no-ILS, CRB – ILS, IBR – ILS, and ABR – ILS, each of which contains 100 simulated datasets.

### Analysis of simulated data

Bayesian divergence times were estimated by using BEAST v2.4.7 (Bouckaert et al., 2014) and RelTime estimates were obtained using MEGA 7 (Kumar, Stecher, & Tamura, 2016; Tamura et al., 2012). Correct substitution model (HKY) and topologies were used in all the dating analyses to prevent confounding errors from phylogeny inference and divergence time estimation.

In BEAST, the root height was calibrated with a normal prior density with mean and standard deviation of 0.05. This was done to exclude the effects of multiple calibrations and the interaction between them, which can impact the results considerably (Battistuzzi, Billing-Ross, Murillo, Filipski, & Kumar, 2015; dos Reis et al., 2018; dos Reis et al., 2015; Duchêne, Duchêne, Holmes, & Ho, 2015). Node ages were inferred based on the Yule (pure birth, BEAST-Yule) and birth-death (BEAST-BD) speciation processes in BEAST, both with default settings, in order to test the performance of distinct Bayesian tree priors. BEAST was also run with coalescent priors, namely the constant population size (BEAST-Constant) and the skyline coalescent (BEAST-Skyline) with default settings (Drummond, Rambaut, Shapiro, & Pybus, 2005). In all analyses, ESS (effective sample sizes) values were checked with the function *effectiveSize* from coda R package (Plummer, Best, Cowles, & Vines, 2006), after discarding the burn-in period.

In RelTime, one does not need to specify a tree prior, a clock model for evolutionary rates, or a statistical distribution to describe the heterogeneity of branch rates (Tamura et al., 2012, 2018). RelTime computes relative times and lineage rates directly from the branch lengths that are inferred from the molecular sequences (Tamura et al., 2012, 2018). Calibrations are also not a prerequisite to estimating divergence times, and so all RelTime inferred times are relative to the height of the ingroup root node.

### Measures

To compare estimated and true times, we report two primary metrics. One is the normalized time difference (Δt), which was computed for each node in every phylogeny analyzed. Δt is the difference between the estimated and the true divergence time divided by the true time. For no-ILS phylogenies, we report Δt for within-population, coalescent, and interspecies comparisons in order to evaluate performance for these three distinct evolutionary levels. However, such a distinction is not possible for ILS datasets. Therefore, we also conducted a linear regression between all the estimated and true times in a phylogeny. The slope of the linear regression through the origin is referred to as the time slope, which is reported alongside the standard R^2^ statistic.

## Results

We carried out a total of 2,400 BEAST analyses for 600 simulated datasets, as four different tree priors were applied for each dataset. The RelTime analysis was conducted only once for each dataset, as RelTime does not require specification of a tree prior. RelTime calculations completed in less than 2 minutes for every dataset, but BEAST analyses took orders of magnitude longer (e.g. Battistuzzi, Billing-Ross, Paliwal, & Kumar, 2011). Also, several BEAST’s MCMC calculations did not reach acceptable ESS values (≥200) even with chain lengths that exceeded one hundred million generations and took ~12 hours to run on an Intel Core i7^®^ iMac @ 4GHz machine. This problem was encountered frequently in BEAST-Yule analyses; they failed for 15% and 32% of no-ILS datasets for IBR and ABR models, respectively. For ILS datasets, 4%, 5% and 3% of ABR replicates in BEAST-Yule, BEAST-BD and BEAST-Skyline did not reach ESS values greater than 200. We excluded these datasets from further consideration and present the results based on the remaining datasets for BEAST analyses.

### Performance for Constant Rate Phylogenies

Biologically, no-ILS phylogenies in which sequence evolved with a constant rate (CBR model) represent the simplest, baseline scenario. For these datasets, distributions of normalized difference between true and estimated time (Δt) for within-population divergences were centered around zero for BEAST (**Fig. 2a**). BEAST-BD and BEAST-Skyline performed the best, and BEAST-Constant showed a slight tendency to overestimate times (**Fig. 2a**). BEAST-Yule overestimated divergence times with much higher median Δt than other tree priors (**Fig. 3a**). In contrast, RelTime showed a tendency to underestimate within-population divergences (median Δt < 0; **Fig. 2a**), because it assigns a time equal to 0 for nodes at which the tip sequences show a small amount of difference.

**Figure 2.**
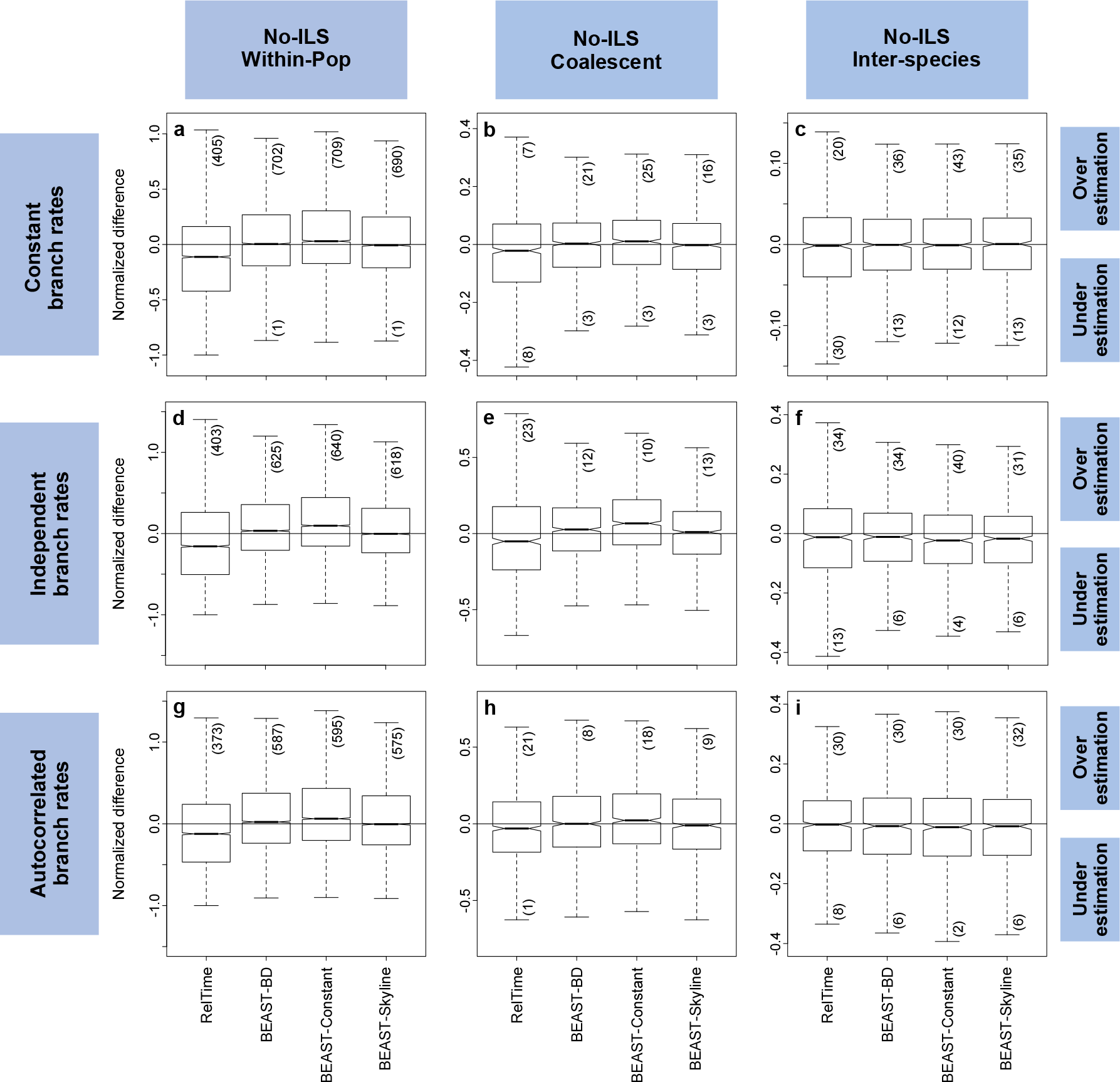
Boxplots of the differences between true and estimated times normalized by the true times (simulations under no-ILS). Constant branch rates in the top panels (**a-c**), independent branch rates on the middle panels (**d-f**), and correlated branch rates on the bottom panels (**g-i**). Left-hand panels **a**, **d** and **g** display normalized difference values for population intra-divergences; center panels **b**, **e** and **h** for species coalescent times; and right-hand panels **c**, **f** and **i** for inter-species divergences. The number inside parentheses indicate the number of outliers, which are not displayed in the figure.

**Figure 3.**
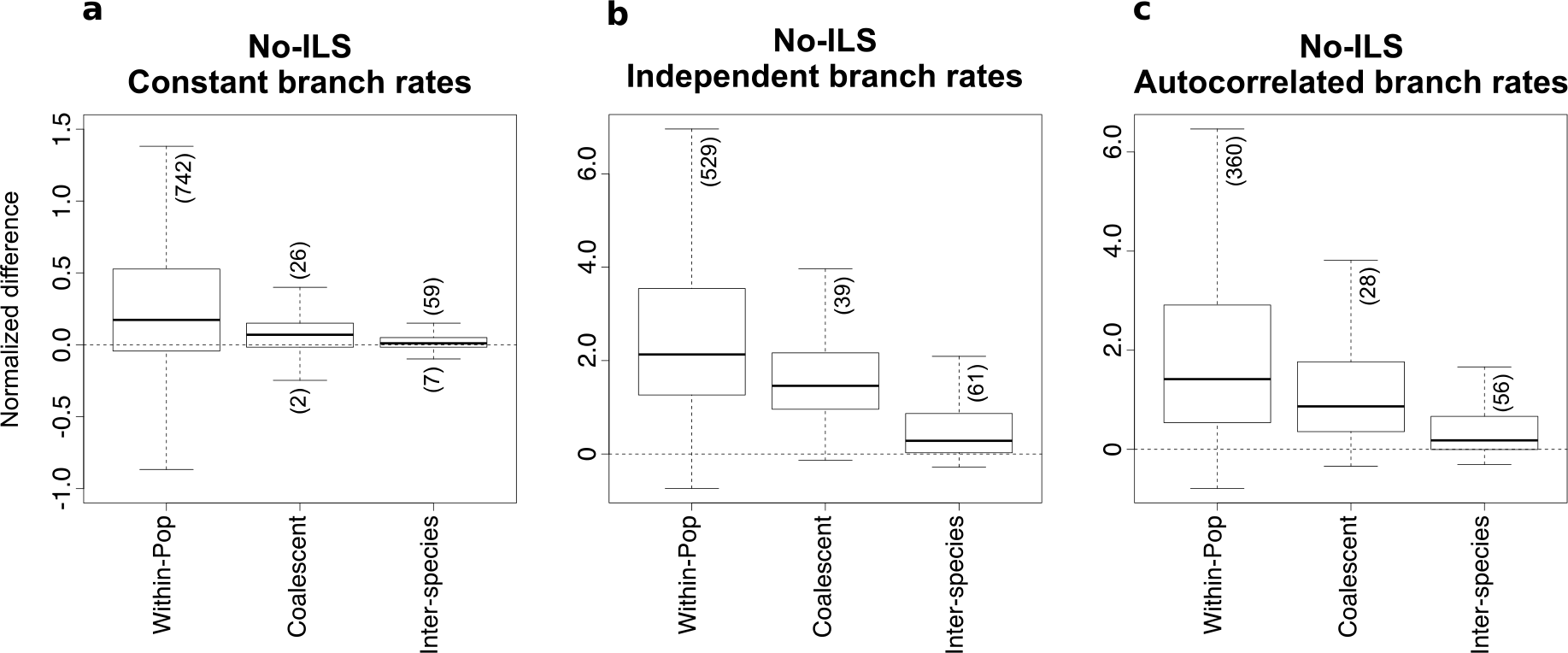
Performance of the Yule prior to estimating divergence times in BEAST under the absence of lineage sorting (simulations under no-ILS). **a-c**, normalized differences between actual and estimated times for intra-population divergences, species tMRCAs, and inter-species divergences. The number inside parentheses indicate the number of outliers, which are not displayed in the figure.

In contrast, BEAST tree priors assign a non-zero time to all the branches (Marin & Hedges, 2018). In our simulations, all the expected divergence times were non-zero, so the difference between the small non-zero divergence times assigned by BEAST and the expected values was smaller than the difference between the zero times assigned by RelTime and the expected values. However, this strategy would result in overestimation of times in BEAST if some of the sampled sequences were truly identical or the sequence divergence was very small. For example, Δt for nodes < 0.1% of the root (ingroup) height was always larger than 0 for BEAST, but it was smaller for RelTime (**Fig. 4**). Consistent with this pattern is the observation that RelTime produced fewer outlier Δt values as compared to BEAST (numbers in parentheses in **Fig. 2a** and **3a**).

**Figure 4:**
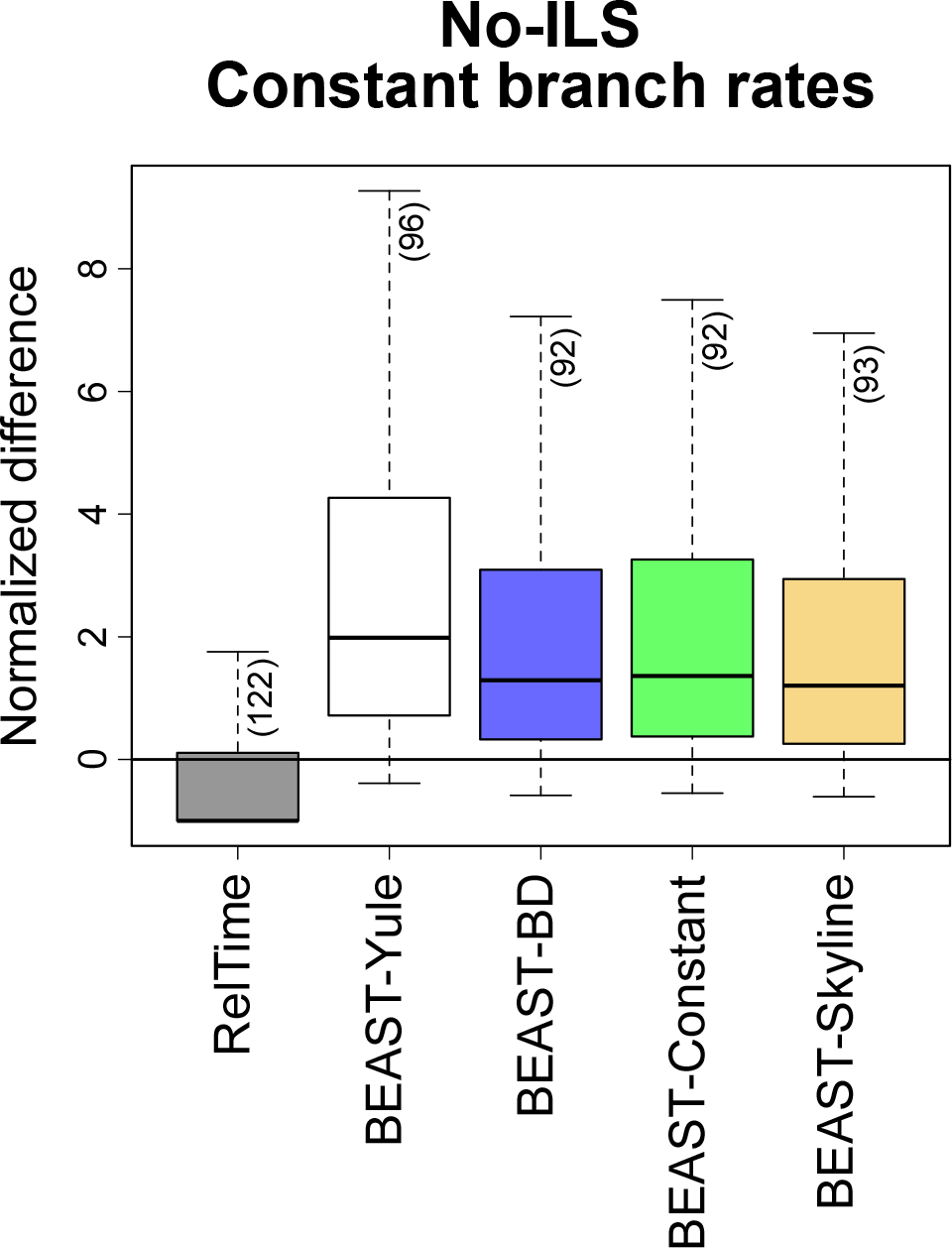
Boxplots of the differences between actual and estimated times normalized by the true times for nodes < 0.1% of the root (ingroup) height in simulations under CBR and no-ILS (100 replicates). The number inside parentheses indicate the number of outliers, which are not displayed in the figure.

Patterns for coalescent time (tMRCA) estimates were similar to those for within-population divergences for BEAST and RelTime (**Fig. 2b** and **3a**), except that the median Δt were closer to zero for BEAST-Constant, BEAST-Yule, and RelTime (**Fig. 2b** and **3a**). RelTime performed better in estimating species tMRCAs than within-population divergences because the effect of very short branches was much smaller. All methods performed well for interspecies comparisons, and Δt values were generally close to zero (**Fig. 2c** and **3a**).

Overall, the analyses of data simulated under a strict clock and without any ILS produced similar patterns among divergence levels. Interestingly, such baseline situations have not been analyzed previously. So, with this baseline, we can begin to assess the impact of biological complexities on divergence time inference over different time scales. For example, a major biological feature of the data utilized in phylogeography studies is the presence of incomplete lineage sorting (ILS). We examined whether the presence of ILS has any impact on node time estimates. For this comparison, we used time slopes, because the presence of ILS does not allow one to distinguish between within-population, coalescence, and inter-species divergences clearly. The distributions of time slopes, derived from 100 simulated datasets in each case, are shown for RelTime (gray), BEAST-Yule (red), BEAST-BD (blue), BEAST-Constant (green), and BEAST-Skyline (yellow) (**Fig. 5**). Generally, BEAST analyses under distinct tree priors and RelTime performed well, with time slopes showing a sharp peak close to 1 for no-ILS (**Fig. 5a**) and ILS phylogenies (**Fig. 5d**). Interestingly, the dispersion of time slopes for ILS phylogenies was smaller than that for the no-ILS phylogenies (**Fig. 5a** and **d**), which is likely because no-ILS phylogenies contained a higher amount of short branches near the tips of the phylogeny as compared to the ILS phylogenies (e.g., **Fig. 1**).

**Figure 5.**
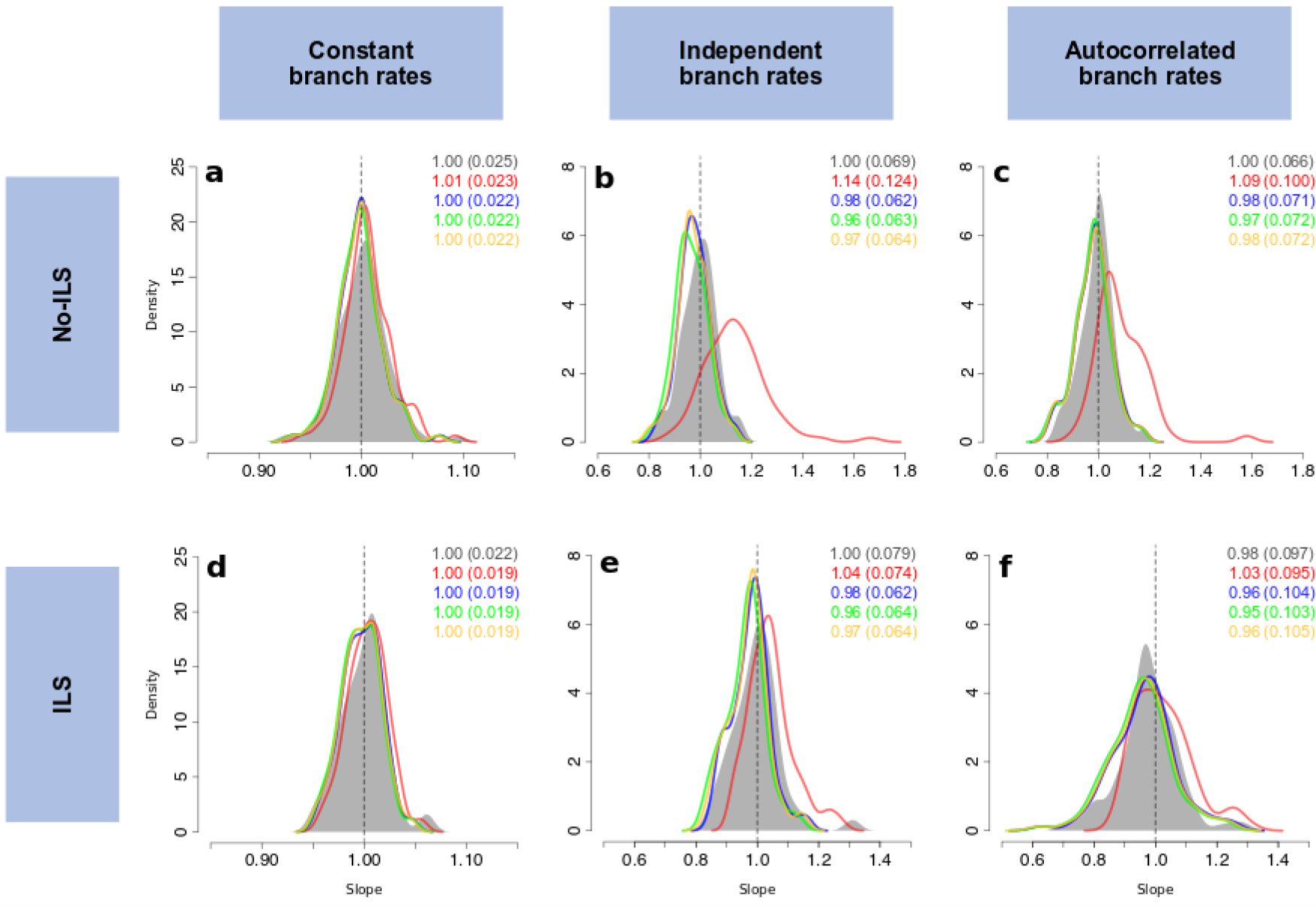
Histogram of slope values resulting from the regression lines between true times and inferred times at RelTime (gray) and BEAST under distinct tree priors: Yule (red), birth-death (blue), constant-size coalescent (green) and skyline coalescent (yellow) (for 100 simulations). Mean and standard deviation (in parentheses) values are displayed inside each panel colored according to the method. Top panels (**a-c**) display results under the absence of lineage sorting (no-ILS), while bottom panels (**d-f**) under the presence of incomplete lineage sorting (ILS). Constant branch rates on the left (**a** and **d**), independent branch rates on the center (**b** and **e**) and correlated branch rates on the right (**c** and **f**).

### Branch Rates Varying Independently (IBR)

The presence of evolutionary rate variability across branches produced results similar to those observed for the strict clock for most of the methods (**Fig. 2** and **4**). The main problem was observed in BEAST-Yule analysis, in which the overestimation of times became more acute (**Fig. 3b**) and time slopes showed clear departure away from 1 (red curves in **Fig. 5b** and **e**). These results agree with Ritchie et al. (2017)’s report about the worse performance of BEAST-Yule for ILS phylogenies, but our analyses showed that this problem becomes even more acute for no-ILS phylogenies (**Fig. 5b**, red curve). RelTime performed well for both no-ILS and ILS phylogenies simulated under the IBR model. For intra-population nodes, RelTime underestimated times slightly (**Fig. 2d**), but in RelTime time slopes were closer to 1 than those from BEAST analyses, although dispersion around the mean was slightly higher for RelTime (**Fig. 5b** and **e**).

As expected, the variability of evolutionary rates caused a greater dispersion of times slopes for BEAST and RelTime for no-ILS datasets (**Fig. 5a** vs. **5b**) and ILS datasets (**Fig. 5d** vs. **5e**). However, the difference in dispersion for no-ILS and ILS datasets was rather small (**Fig. 5a** vs. **5d** and **Fig. 5b** vs. **5e**), which can be observed through the similar standard deviation (SD) values of the distribution of slopes. Overall, we found that all methods performed similarly for no-ILS and ILS phylogenies. The exception was BEAST-Yule analyses, in which the mean of the distribution of slopes was closer to one with SD values considerably lower in ILS simulations (SD=0.124 for no-ILS datasets and SD=0.074 for ILS datasets) (**Fig. 5b** vs. **5e)**.

### Branch rate variation with autocorrelation (ABR)

In the analysis of CBR and IBR datasets, we were able to assume correct evolutionary rate prior in the BEAST analyses. However, BEAST did not have a facility to select an autocorrelated clock model, so we examined how the use of IBR model for ABR datasets impacts performance. There was not much difference between the time slopes for IBR and ABR models in no-ILS phylogenies, except for the BEAST-Yule analyses (**Fig. 5b** vs. **5c**). This means that the violation of the clock model in BEAST when analyzing ABR datasets, caused limited performance detriment. However, the dispersion of time slopes was markedly higher for datasets with ILS (**Fig. 5e** vs. **5f**). RelTime produced a distribution of error that was slightly narrower when compared to BEAST under the BD and coalescent tree priors (**Fig. 5f**). Interestingly, BEAST-Yule’s performance for ABR datasets with ILS turns out to be better than other BEAST analyses (**Fig. 5f**). This result differs from IBR analysis in our simulation and the conclusion of Ritchie et al. (2017), which is made only based on BEAST-Yule’s performance under the IBR model.

## Discussion

Bayesian methods are frequently used to estimate sequence divergence times in inferring phylogeography of species and populations. However, there has been a paucity of computer simulation studies assessing their accuracy, apart from Ritchie’s et al. (2017) study. We have conducted simulations under a variety of scenarios to explore the accuracy of Bayesian time estimates in mixed sample phylogenies with the presence of ILS, the violation of a strict molecular clock, and the misspecification of branch rate model (**Table 1**). Also, we have compared the performance of Bayesian methods with RelTime, a non-Bayesian approach. Thus, we were able to access the impact of tree priors on divergence time estimation in BEAST and showed that BEAST performs well for this kind of dataset. Besides this, we found that RelTime is a reliable and computationally efficient method for estimating divergence times in phylogeographic, phylodynamics and species delimitation studies. In these evaluations, we chose not to study the impact of calibrations on time estimation as other molecular dating studies have done before (Duchêne, Lanfear, & Ho, 2014; Warnock, Parham, Joyce, Lyson, & Donoghue, 2014; Warnock, Yang, & Donoghue, 2017) in order to avoid confounding the impact of calibrations with that of rate models and other priors.

First, we found that RelTime performed well in inferring timetrees for datasets with mixed sampling. RelTime’s performance was comparable to the performance of BEAST when birth-death, constant coalescent and skyline tree priors were used. We found that RelTime’s performance was the least affected by the range of simulated scenarios. It is because RelTime does not require *a priori* specification of tree priors or rate models and directly estimates relative lineage rates and times from branch lengths. In RelTime, the relative rate between sister lineages is the ratio of the evolutionary depths of the two lineages and a relative rate framework is used. This approach contrasts with Bayesian methods, which require the specification of a branch rate model. Therefore, the varying modes of the rate of evolution simulated as well as the presence/absence of ILS did not impact RelTime performance as much as they did impact BEAST under some tree priors. Importantly, RelTime outperformed BEAST in estimating divergences when the branch rates were autocorrelated. As Tao et al. (2019) have shown that the autocorrelation of rates is common in molecular phylogenies, RelTime will perform better than BEAST in empirical data analysis of datasets with a mix of inter and intra-species divergences. RelTime also performed well for ILS phylogenies, which is essential because ILS is widespread for datasets with mixed sampling in phylogeographic studies (Jennings & Edwards, 2005; K. Wang et al., 2018).

We have also found that the performance of BEAST-Yule varied greatly depending on the presence/absence of rate variation, ILS, and rate autocorrelation. Setting the performance of BEAST-Yule aside, one can conclude that the presence of rate variation introduced ≥3x more uncertainty in time estimation (for instance, the SD in slopes’ distribution was ~0.019 and ~0.103 for ILS datasets under CBR and ABR, respectively). The difference was the highest for ILS datasets with the ABR model. Interestingly, our simulation results suggest that the accuracy of dating with datasets containing a mix of inter- and intra-species sampling may not be strongly impacted by the actual shape of the phylogenetic trees, as ILS and no-ILS phylogenies resulted in the similar performance of the dating methods. For BEAST-Yule, the performance was significantly improved when ILS data was analyzed, mainly for ABR simulations. This improvement is likely because of a trade-off between the fact that no-ILS phylogenies contain many more closely-related sequences for which divergence dates are harder to estimate. The number of short branches close to the tips of the phylogeny is much higher in no-ILS than in ILS phylogenies (see **Fig. 1**). This pattern leads to a stronger violation of the Yule process than in the case where ILS was allowed to occur, but it does not appear to impact dating with BD and coalescent priors. This difference is likely because the Yule prior has less flexibility when compared to the other ones, as it considers all divergences to be speciation events and does not model extinction).

Here, for datasets under CBR evolution, the choice of tree priors had little impact on BEAST results. In CBR phylogenies, the confounding effect of times and rates is less problematic than the case in which rates vary among branches. It is, however, important to note that BEAST analyses used a strict clock prior in such cases, so we expect it to perform better than RelTime (which does not assume any *a priori* rate model).

Our analyses confirmed conclusions reached by Ritchie et al. (2017) about the poor performance of BEAST-Yule analyses as compared to BEAST-BD and BEAST-Skyline. Besides this, our analysis of no-ILS phylogenies with IBR model extends this conclusion, as we have found BEAST-Yule to perform even worse for no-ILS datasets as compared to ILS datasets. However, we find that those conclusions do not apply for ABR phylogenies. In the light of the knowledge that the ABR model likely applies for molecular phylogenies and that ILS is likely to be present in mixed sample datasets, BEAST-Yule’s better performance is noteworthy. Overall, however, BEAST-Skyline approach appears to perform the best across all rate and lineage sorting models, which is likely due to the greater flexibility of the Skyline approach (Drummond et al., 2005) that does not assume a constant population size and can accommodate increased diversification in recent times.

Lastly, although equivalent in performance, RelTime generates time estimates orders of magnitude quicker than Bayesian approaches. RelTime has increasingly been used to estimate divergence times for large trees and phylogenomic data (*e.g.* Bond et al. 2014; Mahony et al. 2017; Irisarri et al. 2018; Shin et al. 2018) and was already reported as a robust method for dating large datasets (Filipski, Murillo, Freydenzon, Tamura, & Kumar, 2014; Mello et al., 2017; Tamura et al., 2012). Here, we extend its applicability to datasets that have a mixture of intra- and inter-species sampling, as frequently found in phylogeographic, phylodynamics and species delimitation studies, without assuming *a priori* any model for the branching process on the tree or the branch rate variation.

In conclusion, our simulations covered a wide range of modes of evolution that considered distinct scenarios of rate evolution (CBR, IBR, and ABR) and presence/absence of ILS, many of which was never carried out before for datasets with a mixed sampling of intra- and inter-species sequences. Therefore, our study provides important insights about the impact of tree priors on divergence time estimation in BEAST. Comparisons between BEAST and RelTime’s performance on mixed sampling data also showed that RelTime could be used as a reliable and computationally efficient alternative for estimating divergence times in phylogeographic, phylodynamics and species delimitation studies.

## Acknowledgments

We thank many reviewers for helpful comments on previous versions of this manuscript. This research was supported by grants from the Brazilian Research Council (CNPq, 233920/2014-5, and 409152/2018-8) to BM and National Aeronautics and Space Administration (NASA NNX16AJ30G), National Institutes of Health (GM0126567-02), and National Science Foundation (NSF 1661218) to SK.

## Author contributions

SK and BM conceived the idea, and BM and SK designed research. BM performed analyses; BM, QT, and SK discussed results and wrote the manuscript.

## Data accessibility statement

All the simulated datasets will be made available from FigShare, a publicly accessible repository.

## References

Battistuzzi, F. U., Billing-Ross, P., Murillo, O., Filipski, A., & Kumar, S. (2015). A Protocol for Diagnosing the Effect of Calibration Priors on Posterior Time Estimates: A Case Study for the Cambrian Explosion of Animal Phyla. Molecular Biology and Evolution, 32(7), 1907–1912. doi:10.1093/molbev/msv075

Battistuzzi, F. U., Billing-Ross, P., Paliwal, A., & Kumar, S. (2011). Fast and Slow Implementations of Relaxed-Clock Methods Show Similar Patterns of Accuracy in Estimating Divergence Times. Molecular Biology and Evolution, 28(9), 2439–2442. doi:10.1093/molbev/msr100

Bond, J. E., Garrison, N. L., Hamilton, C. A., Godwin, R. L., Hedin, M., & Agnarsson, I. (2014). Phylogenomics Resolves a Spider Backbone Phylogeny and Rejects a Prevailing Paradigm for Orb Web Evolution. Current Biology, 24(15), 1765–1771. doi:10.1016/j.cub.2014.06.034

Bouckaert, R., Heled, J., Kühnert, D., Vaughan, T., Wu, C.-H., Xie, D., … Drummond, A. J. (2014). BEAST 2: A Software Platform for Bayesian Evolutionary Analysis. PLOS Computational Biology, 10(4), e1003537. doi:10.1371/journal.pcbi.1003537

Chen, W.-C. (2011). Overlapping Codon Model, Phylogenetic Clustering, and Alternative Partial Expectation Conditional Maximization Algorithm. Ph.D. Diss., Iowa Stat University.

Degnan, J. H., & Salter, L. A. (2005). Gene tree distributions under the coalescent process. Evolution; International Journal of Organic Evolution, 59(1), 24–37.

dos Reis, M., Gunnell, G. F., Barba-Montoya, J., Wilkins, A., Yang, Z., & Yoder, A. D. (2018). Using Phylogenomic Data to Explore the Effects of Relaxed Clocks and Calibration Strategies on Divergence Time Estimation: Primates as a Test Case. Systematic Biology, 67(4), 594–615. doi:10.1093/sysbio/syy001

dos Reis, M., & Yang, Z. (2011). Approximate likelihood calculation on a phylogeny for Bayesian estimation of divergence times. Molecular Biology and Evolution, 28(7), 2161–2172. doi:10.1093/molbev/msr045

dos Reis, M., Thawornwattana, Y., Angelis, K., Telford, M. J., Donoghue, P. C. J., & Yang, Z. (2015). Uncertainty in the Timing of Origin of Animals and the Limits of Precision in Molecular Timescales. Current Biology, 25(22), 2939–2950. doi:10.1016/j.cub.2015.09.066

Drummond, A. J., Ho, S. Y. W., Phillips, M. J., & Rambaut, A. (2006). Relaxed Phylogenetics and Dating with Confidence. PLoS Biology, 4(5), e88. doi:10.1371/journal.pbio.0040088

Drummond, A. J., Rambaut, A., Shapiro, B., & Pybus, O. G. (2005). Bayesian Coalescent Inference of Past Population Dynamics from Molecular Sequences. Molecular Biology and Evolution, 22(5), 1185–1192. doi:10.1093/molbev/msi103

Duchêne, D. A., Duchêne, S., Holmes, E. C., & Ho, S. Y. W. (2015). Evaluating the Adequacy of Molecular Clock Models Using Posterior Predictive Simulations. Molecular Biology and Evolution, 32(11), 2986–2995. doi:10.1093/molbev/msv154

Duchêne, S., Lanfear, R., & Ho, S. Y. W. (2014). The impact of calibration and clock-model choice on molecular estimates of divergence times. Molecular Phylogenetics and Evolution, 78, 277–289. doi:10.1016/j.ympev.2014.05.032

Edwards, S. V., & Beerli, P. (2000). Gene divergence, population divergence, and the variance in coalescence time in phylogeographic studies. Evolution; International Journal of Organic Evolution, 54(6), 1839–1854.

Edwards, S. V., Shultz, A. J., & Campbell-Staton, S. C. (2015). Next-generation sequencing and the expanding domain of phylogeography. Folia Zoologica, 64(3), 187–206. doi:10.25225/fozo.v64.i3.a2.2015

Endicott, P., & Ho, S. Y. W. (2008). A Bayesian evaluation of human mitochondrial substitution rates. American Journal of Human Genetics, 82(4), 895–902. doi:10.1016/j.ajhg.2008.01.019

Esselstyn, J. A., Evans, B. J., Sedlock, J. L., Anwarali Khan, F. A., & Heaney, L. R. (2012). Single-locus species delimitation: a test of the mixed Yule-coalescent model, with an empirical application to Philippine round-leaf bats. Proceedings of the Royal Society B: Biological Sciences, 279(1743), 3678–3686. doi:10.1098/rspb.2012.0705

Feller, W. (1939). Die Grundlagen der Volterraschen Theorie Des Kampfes Ums Dasein in Wahrscheinlichkeitstheoretischer Behandlung. Acta Biotheoretica, 5(1), 11–40.

Filipski, A., Murillo, O., Freydenzon, A., Tamura, K., & Kumar, S. (2014). Prospects for Building Large Timetrees Using Molecular Data with Incomplete Gene Coverage among Species. Molecular Biology and Evolution, 31(9), 2542–2550. doi:10.1093/molbev/msu200

Hasegawa, M., Kishino, H., & Yano, T. (1985). Dating of the human-ape splitting by a molecular clock of mitochondrial DNA. Journal of Molecular Evolution, 22(2), 160–174.

Hedges, S. B., Marin, J., Suleski, M., Paymer, M., & Kumar, S. (2015). Tree of life reveals clock-like speciation and diversification. Molecular Biology and Evolution, 32(4), 835–845. doi:10.1093/molbev/msv037

Heled, J., & Drummond, A. J. (2010). Bayesian Inference of Species Trees from Multilocus Data. Molecular Biology and Evolution, 27(3), 570–580. doi:10.1093/molbev/msp274

Ho, S. Y. W. (2014). The changing face of the molecular evolutionary clock. Trends in Ecology and Evolution, 29(9), 496–503. doi:10.1016/j.tree.2014.07.004

Ho, S. Y. W., Duchêne, S., & Duchêne, D. A. (2015). Simulating and detecting autocorrelation of molecular evolutionary rates among lineages. Molecular Ecology Resources, 15(4), 688–696. doi:10.1111/1755-0998.12320

Hudson, R. R. (2002). Generating samples under a Wright-Fisher neutral model of genetic variation. Bioinformatics, 18(2), 337–338. doi:10.1093/bioinformatics/18.2.337

Irisarri, I., Singh, P., Koblmüller, S., Torres-Dowdall, J., Henning, F., Franchini, P., … Meyer, A. (2018). Phylogenomics uncovers early hybridization and adaptive loci shaping the radiation of Lake Tanganyika cichlid fishes. Nature Communications, 9(1), 3159. doi:10.1038/s41467-018-05479-9

Jennings, W. B., & Edwards, S. V. (2005). Speciational history of Australian grass finches (Poephila) inferred from thirty gene trees. Evolution; International Journal of Organic Evolution, 59(9), 2033–2047.

Kendall, D. G. (1948). On the Generalized “Birth-and-Death” Process. The Annals of Mathematical Statistics, 19(1), 1–15.

Kishino, H., Thorne, J. L., & Bruno, W. J. (2001). Performance of a divergence time estimation method under a probabilistic model of rate evolution. Mol Biol Evol, 18(3), 352–361. doi:10.1093/oxfordjournals.molbev.a003811

Kuhner, M. K., Yamato, J., & Felsenstein, J. (1995). Estimating effective population size and mutation rate from sequence data using Metropolis-Hastings sampling. Genetics, 140(4), 1421–1430.

Kuhner, M. K., Yamato, J., & Felsenstein, J. (1998). Maximum Likelihood Estimation of Population Growth Rates Based on the Coalescent. Genetics, 149(1), 429–434.

Kumar, S., Stecher, G., Li, M., Knyaz, C., & Tamura, K. (2018). MEGA X: Molecular Evolutionary Genetics Analysis across Computing Platforms. Molecular Biology and Evolution, 35(6), 1547–1549. doi:10.1093/molbev/msy096

Kumar, S., Stecher, G., & Tamura, K. (2016). MEGA7: Molecular Evolutionary Genetics Analysis Version 7.0 for Bigger Datasets. Molecular Biology and Evolution, 33(7), 1870–1874. doi:10.1093/molbev/msw054

Lemmon, A. R., & Lemmon, E. M. (2012). High-Throughput Identification of Informative Nuclear Loci for Shallow-Scale Phylogenetics and Phylogeography. Systematic Biology, 61(5), 745–761. doi:10.1093/sysbio/sys051

Liu, L., & Yu, L. (2010). Phybase: An R package for species tree analysis. Bioinformatics, 26(7), 962–963. doi:10.1093/bioinformatics/btq062

Maddison, W. P. (1997). Gene Trees in Species Trees. Systematic Biology, 46(3), 523–536. doi:10.1093/sysbio/46.3.523

Mahony, S., Foley, N. M., Biju, S. D., & Teeling, E. C. (2017). Evolutionary History of the Asian Horned Frogs (Megophryinae): Integrative Approaches to Timetree Dating in the Absence of a Fossil Record. Molecular Biology and Evolution, 34(3), 744–771. doi:10.1093/molbev/msw267

Manzanilla, V., Kool, A., Nguyen Nhat, L., Nong Van, H., Le Thi Thu, H., & de Boer, H. J. (2018). Phylogenomics and barcoding of Panax: toward the identification of ginseng species. BMC Evolutionary Biology, 18, 44. doi:10.1186/s12862-018-1160-y

Marin, J., & Hedges, S. B. (2018). Undersampling Genomes has Biased Time and Rate Estimates Throughout the Tree of Life. Molecular Biology and Evolution, 35(8), 2077–2084. doi:10.1093/molbev/msy103

McCluskey, B. M., & Postlethwait, J. H. (2015). Phylogeny of Zebrafish, a “Model Species,” within Danio, a “Model Genus.” Molecular Biology and Evolution, 32(3), 635–652. doi:10.1093/molbev/msu325

McCormack, J. E., Hird, S. M., Zellmer, A. J., Carstens, B. C., & Brumfield, R. T. (2013). Applications of next-generation sequencing to phylogeography and phylogenetics. Molecular Phylogenetics and Evolution, 66(2), 526–538. doi:10.1016/j.ympev.2011.12.007

Mello, B., Tao, Q., Tamura, K., & Kumar, S. (2017). Fast and Accurate Estimates of Divergence Times from Big Data. Molecular Biology and Evolution, 34(1), 45–50. doi:10.1093/molbev/msw247

Mello, B., Vilela, J. F., & Schrago, C. G. (2018). Conservation phylogenetics and computational species delimitation of Neotropical primates. Biological Conservation, 217, 397–406. doi:10.1016/j.biocon.2017.11.017

Melville, J., Haines, M. L., Boysen, K., Hodkinson, L., Kilian, A., Smith Date, K. L., … Parris, K. M. (2017). Identifying hybridization and admixture using SNPs: application of the DArTseq platform in phylogeographic research on vertebrates. Royal Society Open Science, 4(7), 161061. doi:10.1098/rsos.161061

Merwe, M., McPherson, H., Siow, J., & Rossetto, M. (2014). Next-Gen phylogeography of rainforest trees: exploring landscape-level cpDNA variation from whole-genome sequencing. Molecular Ecology Resources, 14(1), 199–208. doi:10.1111/1755-0998.12176

Ogilvie, H. A., Bouckaert, R. R., & Drummond, A. J. (2017). StarBEAST2 Brings Faster Species Tree Inference and Accurate Estimates of Substitution Rates. Molecular Biology and Evolution, 34(8), 2101–2114. doi:10.1093/molbev/msx126

Ogilvie, H. A., Heled, J., Xie, D., & Drummond, A. J. (2016). Computational Performance and Statistical Accuracy of *BEAST and Comparisons with Other Methods. Systematic Biology, 65(3), 381–396. doi:10.1093/sysbio/syv118

O’Reilly, J. E., dos Reis, M., & Donoghue, P. C. J. (2015). Dating Tips for Divergence-Time Estimation. Trends in Genetics, 31(11), 637–650. doi:10.1016/j.tig.2015.08.001

Plummer, M., Best, N., Cowles, K., & Vines, K. (2006). CODA: convergence diagnosis and output analysis for MCMC. R News, 6(1), 7–11.

Rambaut, A., & Grass, N. C. (1997). Seq-Gen: an application for the Monte Carlo simulation of DNA sequence evolution along phylogenetic trees. Comput Appl Biosci, 13(3), 235–238. doi:10.1093/bioinformatics/13.3.235

Rannala, B., & Yang, Z. (2003). Bayes estimation of species divergence times and ancestral population sizes using DNA sequences from multiple loci. Genetics, 164(4), 1645–1656.

Rannala, B., & Yang, Z. (2017). Efficient Bayesian Species Tree Inference under the Multispecies Coalescent. Systematic Biology, 66(5), 823–842. doi:10.1093/sysbio/syw119

Rosenberg, M. S., & Kumar, S. (2003). Heterogeneity of nucleotide frequencies among evolutionary lineages and phylogenetic inference. Molecular Biology and Evolution, 20(4), 610–621. doi:10.1093/molbev/msg067

Shin, S., Clarke, D. J., Lemmon, A. R., Moriarty Lemmon, E., Aitken, A. L., Haddad, S., … McKenna, D. D. (2018). Phylogenomic Data Yield New and Robust Insights into the Phylogeny and Evolution of Weevils. Molecular Biology and Evolution, 35(4), 823–836. doi:10.1093/molbev/msx324

Stadler, T. (2011). Simulating trees with a fixed number of extant species. Systematic Biology, 60(5), 676–684. doi:10.1093/sysbio/syr029

Takahata, N. (1989). Gene genealogy in three related populations: consistency probability between gene and population trees. Genetics, 122(4), 957–966.

Takahata, N., & Nei, M. (1985). Gene genealogy and variance of interpopulational nucleotide differences. Genetics, 110(2), 325–344.

Tamura, K., Battistuzzi, F. U., Billing-Ross, P., Murillo, O., Filipski, A., & Kumar, S. (2012). Estimating divergence times in large molecular phylogenies. Proceedings of the National Academy of Sciences, 109(47), 19333–19338. doi:10.1073/pnas.1213199109

Tamura, K., Tao, Q., & Kumar, S. (2018). Theoretical Foundation of the RelTime Method for Estimating Divergence Times from Variable Evolutionary Rates. Molecular Biology and Evolution, 35(7), 1770–1782. doi:10.1093/molbev/msy044

Tao, Q., Tamura, K., Battistuzzi, F., & Kumar, S. (2019). A new method for detecting autocorrelation of evolutionary rates in large phylogenies. Molecular Biology and Evolution. doi:10.1093/molbev/msz014

Wang, C., Shikano, T., Persat, H., & Merilä, J. (2015). Mitochondrial phylogeography and cryptic divergence in the stickleback genus Pungitius. Journal of Biogeography, 42(12), 2334–2348. doi:10.1111/jbi.12591

Wang, K., Lenstra, J. A., Liu, L., Hu, Q., Ma, T., Qiu, Q., & Liu, J. (2018). Incomplete lineage sorting rather than hybridization explains the inconsistent phylogeny of the wisent. Communications Biology, 1, 169. doi:10.1038/s42003-018-0176-6

Warnock, R. C. M., Parham, J. F., Joyce, W. G., Lyson, T. R., & Donoghue, P. C. J. (2014). Calibration uncertainty in molecular dating analyses: there is no substitute for the prior evaluation of time priors. Proceedings of the Royal Society B: Biological Sciences, 282(1798), 20141013–20141013. doi:10.1098/rspb.2014.1013

Warnock, R. C. M., Yang, Z., & Donoghue, P. C. J. (2017). Testing the molecular clock using mechanistic models of fossil preservation and molecular evolution. Proc. R. Soc. B, 284(1857), 20170227. doi:10.1098/rspb.2017.0227

Xu, B., & Yang, Z. (2016). Challenges in Species Tree Estimation Under the Multispecies Coalescent Model. Genetics, 204(4), 1353–1368. doi:10.1534/genetics.116.190173

Yang, Z. (2015). The BPP program for species tree estimation and species delimitation. Current Zoology, 61(5), 854–865. doi:10.1093/czoolo/61.5.854

Yang, Z., & Rannala, B. (2014). Unguided Species Delimitation Using DNA Sequence Data from Multiple Loci. Molecular Biology and Evolution, 31(12), 3125–3135. doi:10.1093/molbev/msu279

Yule, G. U. (1924). A mathematical theory of evolution, based on the conclusions of Dr. J. C. Willis. Phil. Trans. R. Soc. Lond. B, 213, 21–87. doi:10.1098/rstb.1925.0002

